# Efficient data management infrastructure for the integration of imaging and omics data in life science research

**DOI:** 10.1101/2019.12.28.889295

**Authors:** Luis Kuhn Cuellar, Andreas Friedrich, Gisela Gabernet, Luis de la Garza, Sven Fillinger, Adrian Seyboldt, Sven zur Oven-Krockhaus, Friederike Wanke, Sandra Richter, Wolfgang M. Thaiss, Marius Horger, Nisar Malek, Klaus Harter, Michael Bitzer, Sven Nahnsen

**Affiliations:** Quantitative Biology Center (QBiC), University of Tübingen, Tübingen, Germany; Center for Plant Molecular Biology (ZMBP), University of Tübingen, Tübingen, Germany; Department of Radiology, Diagnostic and Interventional Radiology, University of Tübingen, Tübingen, Germany; Department Internal Medicine I, University of Tübingen, Tübingen, Germany

## Abstract

As technical developments in omics and biomedical imaging drive the increase in quality, modality, and throughput of data generation in life sciences, the need for information systems capable of integrative, long-term storage and management of these heterogeneous digital assets is also increasing. Here, we propose an approach based on principles of Service Oriented Architecture design, to allow the integrated management and analysis of multi-omics and biomedical imaging data. The proposed architecture introduces an interoperable image management system, the OMERO server, into the backend of qPortal, a FAIR-compliant web-based platform for omics data management. The implementation of an integrated metadata model, the development of software components to enable the communication with the OMERO server, and an extension to the data management operations of established software, allows for FAIR management of heterogeneous omics and biomedical imaging data within an integrated system, which facilitates metadata queries from web-based scientific applications. The applicability of the proposed architecture is demonstrated in two prototypical use cases, a plant biology study using confocal scanning microscopy, and a clinical study on hepatocellular carcinoma, with data from a variety of medical imaging and omics modalities. We anticipate that FAIR data management systems for multi-modal data repositories will play a pivotal role in data-driven research, as the joint analysis of omics and imaging data becomes not only desirable but necessary to derive novel insights into biological processes. In particular for powerful machine learning applications where the availability of large datasets with high quality phenotypic annotations is a requirement.

## Introduction

Current technical advances in the life sciences allow researchers to collect large amounts of data that open fundamentally new routes for a multitude of research questions. Typically, multi-omics and biomedical imaging data from biological samples are building the most important data basis in such studies. The large volume of data produced by various omics disciplines (e.g. genomics, transcriptomics, proteomics, metabolomics), biomedical imaging techniques (e.g. X-ray CT and PET), and the unprecedented increase in spatial resolution achieved by conventional confocal and super-resolution light microscopy (Sigal, Zhou, & Zhuang, 2018) and modern electron microscopes (Cheng, 2015), present a challenge for long-term storage and efficient management of these high-dimensional digital assets, especially with regard to the requirements imposed by the FAIR data management principles (Wilkinson et al., 2016). Moreover, it is of particular importance to employ rich metadata models that allow researchers to relate data from different disciplines, and design experiments using an integrative approach to handle both, multilayer omics, as well as biomedical imaging data. The increasing data volume and experimental design complexity call for data management systems allowing for the integration of multi-omics and imaging data.

In recent years, a variety of data management and analysis systems for life science have been introduced. Among many, two examples are the Galaxy platform, a web-based workflow system for reproducible genomic analysis (Goecks et al. 2010; Afgan et al. 2016), and the cBio Portal, which focuses on exploration and visualization of large datasets of cancer genomics data (Cerami et al. 2012). For proteomics studies, the Swiss Grid Proteomics Portal (iPortal) provides web-based analysis tools (Kunszt et al. 2015). The Open Microscopy Environment Remote Objects (OMERO) is a sophisticated image management system for biology and medicine. Despite its popularity, it offers limited support for managing omics data (Allan et al. 2012). While several systems focus on the management of omics or imaging data (Bauch et al. 2011; Allan et al. 2012), or are dedicated to bioinformatics workflows (Goecks et al. 2010; Afgan et al. 2016; Chaumont et al. 2012), to our knowledge no approach has been suggested to provide a solution for the integrated management of multi-omics and imaging data in parallel. Additionally, most of the available platforms focus on a limited set of omics disciplines, thus leading to metadata models that are not well-suited to describe complex experimental designs with multiple omics and imaging modalities.

The lack of comprehensive and validated metadata storage is one of the pitfalls of reproducible research in the genomic age (Baker 2016). Accordingly, many researchers are starting to embrace proposed standards, such as the FAIR data principles, for omics data management. To address the aforementioned issues, we recently introduced qPortal, a web-based solution, which follows FAIR guidelines, for the management of the entire added-value chain of omics-based biomedical data (Mohr et al. 2018). qPortal is a web-based portal for scientific data management and analysis, which uses the Open Biological Information System (openBIS) platform (Bauch et al., 2011) as a backend. While FAIR data management for genomics and proteomics studies has been well established using qPortal, it lacks the capability of managing imaging data (e.g. from light microscopy or MRI).

Here, we propose a solution based on principles of Service Oriented Architecture (SOA) design (Josuttis 2007), to integrate an interoperable image management system for the accommodation of data from microscopy and medical imaging studies in qPortal. Our suggested infrastructure enables the integrated analysis of imaging and omics data, which is an increasingly deployed strategy in biology and medical studies (Stoyanova et al. 2016; Disselhorst et al. 2018; Hériché et al. 2019). We suggest using an OMERO server as a backend component for all imaging data. In architectural conjunction with openBIS, building the conventional backbone for qPortal. The OMERO server is capable of enforcing FAIR principles for the imaging data itself, while providing large-scale data storage, management, and compatibility with common imaging file formats (Linkert et al., 2010).On the other hand, the openBIS server ensures FAIR principles on the metadata describing the experimental design of research projects and the resulting omics data, while qPortal provides a web-based interface to present users with a set of applications for scientific data management.

## Methodology

Our implementation efficiently leverages the openBIS, qPortal, and OMERO platforms to provide FAIR data management for multi-modal data in life science research. Specifically, it allows management and analysis of omics data (e.g. gene sequencing and expression data, mass spectrometry) in conjunction with data from microscopy and medical imaging disciplines. Integration of omics and imaging data modalities at the metadata level is achieved by a combination of well-established metadata models (i.e. the qPortal and OMERO models), and implemented through middleware software that provides the logic, and connective tissue between the application program interface (API) of the aforementioned platforms, allowing them to work in concert as components of a unified system.

### Architecture

Our data management architecture integrates the OMERO platform into qPortal, it leverages openBIS and OMERO functionality to facilitate the integration of both imaging and omics data within qPortal. openBIS serves the purpose of storing and managing raw data and metadata from omics sources, including the general experimental design of research projects and basic information regarding the biology of the samples. On the other hand, OMERO is best suited for modeling, storing and managing medical imaging and microscopy data, employing a metadata model focused on describing multi-channel spatio-temporal data, the technical specification of imaging instruments, and the experimental parameters used during data acquisition.

While the openBIS and OMERO servers function as the backend of the system, qPortal provides a web-based user interface with a set of applications for scientific data management. These applications support various project management operations and orchestrate the complex data management tasks by using utility components designed to facilitate a connection to the openBIS and OMERO servers for metadata transfer. **Fig. 1** depicts the component diagram of the proposed architecture.

**Fig. 1.**
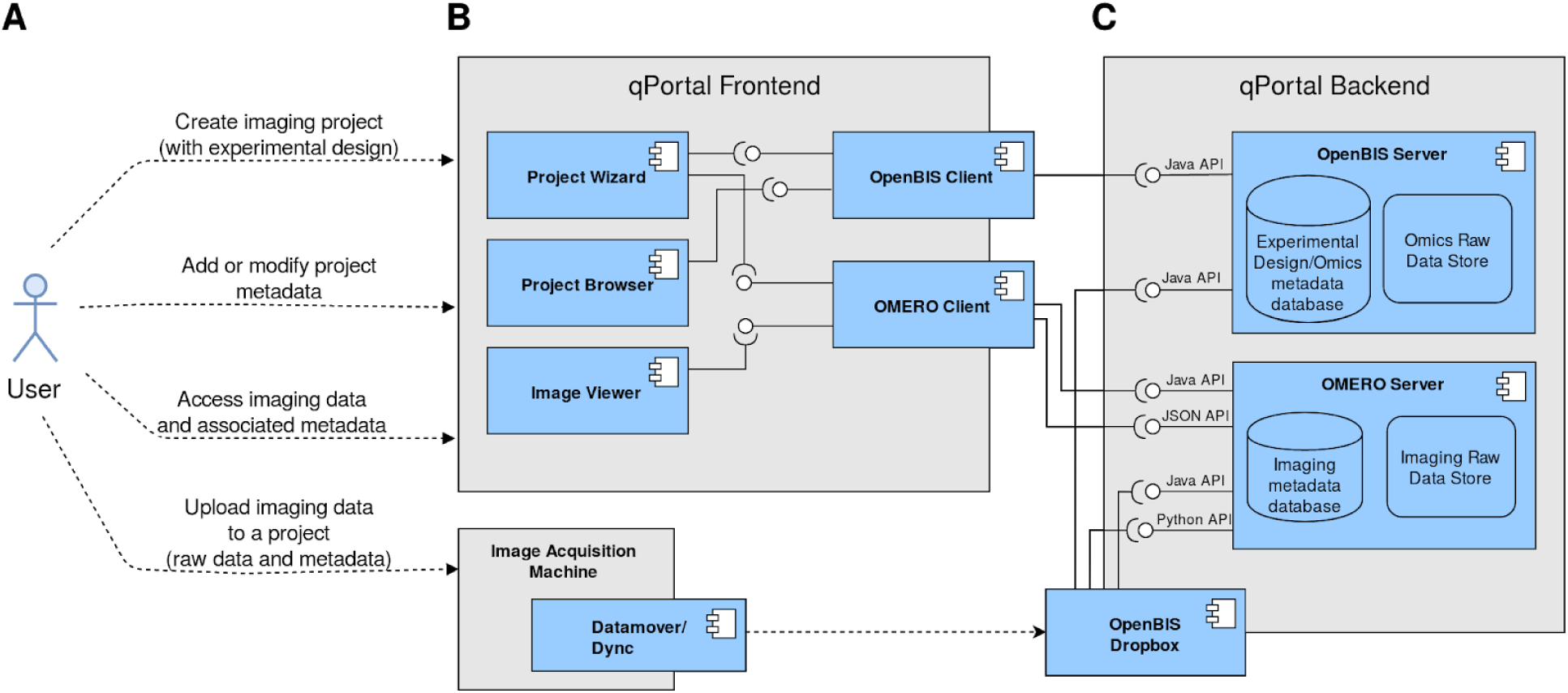
Component diagram of the proposed data management infrastructure for omics and imaging data, indicating the interaction of all major software components. **(A)** Users can perform a set of data management operations, including uploading new imaging data into a previously created project, by copying it into a dropbox folder. **(B)** The qPortal platform as a frontend, containing portlet applications that connect to the backend via connector components (openBIS and OMERO clients). **(C)** The backend of the proposed system comprises two major components, the openBIS and OMERO servers.

Raw imaging data and accompanying metadata can be uploaded into the system using an openBIS-based *dropbox* mechanism. In short, data is securely transferred to an *incoming server* within the backend of the proposed system, a process is then triggered to allow for structured ingestion of raw data and metadata. This process uploads images to the OMERO server and stores the resulting unique identifiers in the openBIS server as metadata items, in order to maintain a consistent record in both servers.

### The OMERO Platform

The OME Remote Objects (OMERO) is a data management platform for biological imaging that is based on the OME Data Model (Goldberg et al., 2005) and the Bio-Formats project (Linkert et al., 2010). It is composed of databases, middleware and remote client applications. The main component is the OMERO.server, a Java application that acts as middleware connecting various databases that store different data types, and provides access to stored data via a single API. The OMERO.server not only provides access to the underlying storage facilities, it must also process data before delivering it to the client applications.

OMERO provides a Java-based client for imaging data upload, the OMERO.importer. This tool is a command line interface that uses Bio-Formats to read image data and metadata from supported file formats. During data import, Bio-Formats reads metadata from the raw data file and maps it to the OME Data Model (Goldberg et al., 2005). While a binary repository is used to store binary data such as images and thumbnails, the OMERO relational database archives all metadata (see metadata models) associated with binary images, user information, and simple data annotations.

Since modern biological images may consist of various image frames recorded at different positions, channels, or timepoints, the OMERO.server contains a multi-threaded image rendering engine that can rapidly display image planes, and transfer them via the API to clients. This engine reads images from the binary repository and can apply transforms, according to the parameters provided by an OMERO client or the OMERO relational database. Supported operations include image compression, overlay and projection. Users can therefore create multiple views of the data without modifying the originally acquired data.

While data can be imported using the provided OMERO.importer or OMERO.dropbox tools (a filesystem monitoring tool), third-parties can also develop specialized tools to access metadata and raw imaging data by querying the Java or Python APIs of the OMERO.server. Remote data access between client applications and the OMERO.server is achieved via ZeroC’s Internet Communication Engine (Henning 2004).

OMERO also includes a customizable web-based client for image visualization and annotation. OMERO.web is a Django-based application that uses the APIs of the OMERO.server to provide a web interface for metadata access, metadata annotation, and full 5D (i.e. space, time, and channels) image visualization.

Regions of interest (ROI) describing the spatial boundaries of detected objects in an image, which are often the product of manual annotation or segmentation algorithms, can be stored in the OMERO database as geometric objects (e.g. points, circles, polygons), thus supporting ROI storage for 5D images.

### The OpenBIS platform

The openBIS platform is an open source management system for data acquired in biological experiments (Bauch et al. 2011). The main components of this platform are a data store, a flexible and extendable metadata model supporting complex XML-based annotation, a relational database to store metadata, and an application server to browse, access and manage both data and metadata.

In order to upload data into openBIS, a *dropbox* is usually employed. In this *context*, a *dropbox* is a filesystem monitoring mechanism that can transfer files containing raw data and metadata, from an acquisition machine or adjacent computer within a source laboratory, to a server in the backend of the data management system, which temporarily stores *incoming* data. Usually, this data transfer operation is carried out securely using the openBIS Datamover or the Dync application (Seyboldt and Fillinger, 2019).

Subsequently, this *incoming* backend server executes the *extract, transform, load* (ETL) routine associated with the specific source laboratory and raw data type, the objective of this routine is to process raw data before uploading it into the openBIS data store. ETL routines can execute additional external scripts and often extract metadata from raw data files. During the ETL process all the necessary metadata entities (e.g. experiments, samples, datasets) are created, and all available metadata is stored as properties of the newly created entities.

### The Web-based *qPortal*

qPortal is a web-based platform for FAIR data management of biomedical data. It enforces a well-structured metadata model to capture the experimental design of projects and the biology of the samples. This platform is built on a Liferay instance and contains a large collection of loosely coupled portlets (i.e. java-based, web applications) that use the open-source framework VAADIN (Duarte, 2013). qPortal also offers integrated workflow support on stored data. The main portlet applications, the *Project Wizard* and *Project Browser*, facilitate the creation of projects with complex experimental designs, and provide data access and management, respectively.

The registration of an experimental design is facilitated with the *Project Wizard*. The application helps users to register a new project by taking them through a series of defined steps to describe the experimental design and metadata of the project. The first step describes the biological entities under study (e.g. patients, model organism) on species level, the second step captures the extraction of cells or tissues from biological entities. Finally, the third step describes the process used to prepare the biological sample for data acquisition. Experimental factors such as treatment or genotype can also be defined precisely and in order to maximize statistical power a full-factorial design is proposed by default. Once all this information has been collected, the *Project Wizard* creates and stores all the metadata entities and relations in the openBIS database, according to a well-defined metadata model (see metadata model section).

The main interaction with project-associated data is facilitated by the *Project Browser*. This application allows search and access to project metadata, as well as providing direct access to raw data and analysis results. This portlet shows all projects the logged-in user has access to, in a searchable table that provides general project information, e.g. project code and description. By clicking on a project, the user can access all project metadata, which the application visualizes as a tree-like structure of metadata entities (**Fig. S1A** and metadata models). Metadata concerning the biology of the samples can be accessed, and the associated raw data can be directly downloaded. Data resulting from workflow executions can also be accessed from a dedicated section within the application.

As part of the qPortal backend, an openBIS server instance provides the means of storing and managing raw data and metadata. While the *Project Wizard* application only registers the main information of the experiments and project, the openBIS-based ETL routines register the remainder of the experimental metadata concerning the file (e.g. sample preparation and data acquisition parameters) during data upload. Effectively, the *Project Wizard* and ETL scripts enforce the FAIR-compliant metadata structure in a consistent manner for all projects.

qPortal provides an interface to allow execution of computations in cluster infrastructures using the data stored in openBIS, facilitating the automation of common bioinformatics analyses. Once a workflow has been executed, qPortal can register the resulting data into openBIS with a metadata reference to the original input data and the parameters used during analysis, to allow reproducibility of workflow results. Moreover, workflow systems such as gUSE (Kacsuk et al. 2012) and Nextflow (Di Tommaso et al., 2017) have been successfully connected to the qPortal platform. In particular, qPoral is compatible with the nf-core framework (Ewels et al. 2020), which provides access to standardized and reproducible bioinformatics pipelines, along with the necessary pipeline development tools.

## Results

### Integration of OMERO into the qPortal Platform

The integration of an OMERO server as an additional backend component of qPortal, requires substantial changes in the data handling and managing processes, considering that metadata information between the openBIS and OMERO servers has to be connected and kept synchronized in a structured manner (**Fig. S1**). First, an integrative model was established, which defines detailed metadata boundaries between the project and omics domains, as managed by the openBIS server, and the imaging domain stored in the OMERO server. The technical implementation of this model requires extending server-side qPortal applications and processes, and the development of modules to facilitate communication between these applications and the OMERO server.

In order to allow communication between the OMERO server and qPortal, we developed a Java library wrapping the OMERO server API. This *OMERO client* library is analog to the *openBIS Client* component of qPortal (Mohr et al., 2018), and is used by the *Project Wizard* application to implement the metadata model connection during project creation, i.e. creating synchronized metadata entities and structures in both the openBIS and OMERO platforms (see metadata models).

To effectively provide web-based image visualization in qPortal, we developed an *Image Viewer* portlet. This application provides easy access to imaging metadata and image visualization, using the *OMERO Client* component to query the OMERO server. The *Image Viewer* application also accesses OMERO.web functionality, in particular the 5D image viewers.

Finally, a suitable ETL routine for imaging data was created. This routine uploads image data into the OMERO platform, and creates a record of the available imaging data in the openBIS server. This can be achieved by creating metadata entities in the openBIS database that contain symbolic links to the images stored in OMERO binary repository (see **Fig. 2**).

**Fig. 2.**
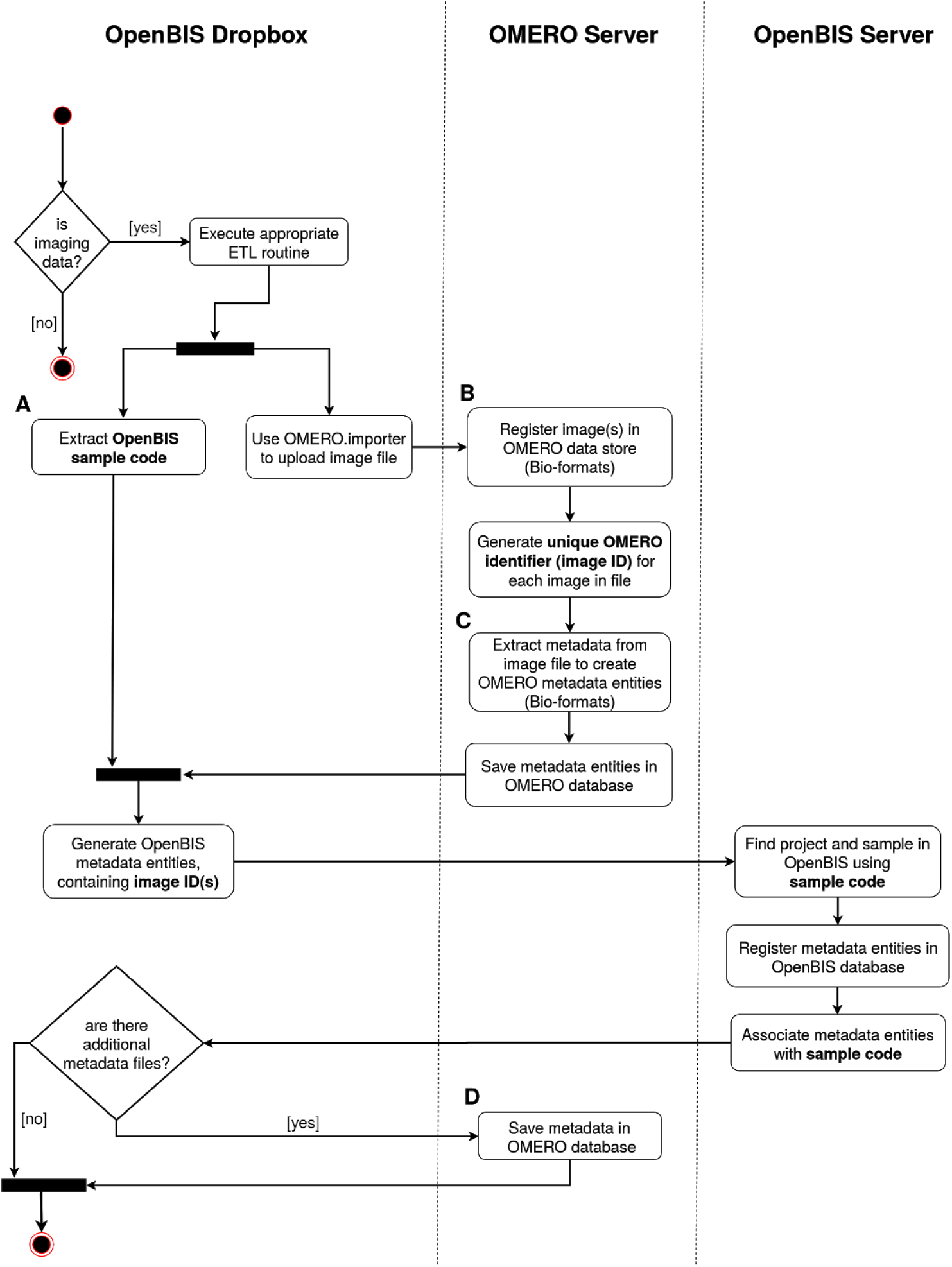
Activity diagram of the ETL process for imaging data. The diagram indicates which tasks are executed on each of the system components involved in the image registration process. **(A)** The openBIS sample code in this step was previously generated by the *Project Wizard*. **(B)** Image files are parsed by Bio-formats, providing compatibility with a large collection of imaging file formats. **(C)** Bio-formats extracts metadata from image files and maps it to the OMERO metadata model. **(D)** Save tabular metadata as key-value pairs associated with the previously registered images.

#### Metadata Models

The full data management potential of both OMERO and qPortal can be achieved within a unified system, when their metadata models are centralized, or at least connected in a systematic way. This allows for full access to the metadata management capacities of openBIS, the metadata structure for project and multi-factorial experiments implemented by qPortal, including its domain-specific and feature-rich metadata support, while using the metadata model and storage facilities of OMERO to describe microscopy and medical imaging data and its acquisition process. It is imperative that both models are linked so that any metadata transaction preserves the correct execution of CRUD operations (create, read, update, delete) on the semantic level, considering the databases of both platforms. Metadata redundancies, and ambiguous, or orphaned metadata relations have to be avoided. Since the metadata model of both platforms has a hierarchical organization, with conceptually similar metadata entities in the main structure, it is straightforward to align and connect metadata entities across models using cardinality relationships (e.g. one-to-many, many-to-many). The general openBIS model is composed of four main hierarchical levels, with project entities on top, followed by experiments, samples, and finally datasets. Similarly, the OMERO model uses a hierarchical structure, with project entities on top, followed by datasets, images, pixels, and finally features. In order to connect both metadata structures, we propose a loose-coupling approach, following SOA principles (Josuttis 2007). Here, two major one-to-one correspondences are established, one between “project” entities and another one relating qPortal sample entities with OMERO dataset entities. **Fig. S1** depicts the unified metadata model.

While both models can handle extensible XML schemas, we aim to distribute metadata information in a manner that benefits from the most useful features of each platform. Therefore, we allow the qPortal model to continue describing the main experimental design of the project and capture the biology of the samples, while using the OMERO model to store image related metadata, e.g. pixel size, magnification, image acquisition parameters, technical specifications of the microscope or medical imaging device. With this approach we can benefit from both Bio-Formats and the project planning functionality offered by qPortal.

#### Technical Implementation

The *OMERO client* module was implemented as a java-based library (https://github.com/qbicsoftware/omero-client-lib). It provides an interface to read and write metadata on the OMERO server from portlet applications, the interface is tailored to implement the necessary operations needed to maintain the proposed coupling of metadata models. This library communicates with the OMERO server via the OMERO Java API and encapsulates connectivity logic with the backend, allowing portlet applications to focus on user interface and metadata synchronization between the openBIS and OMERO servers.

The *Project Wizard* application (https://github.com/qbicsoftware/projectwizard-portlet) was extended to provide the user with an option to enable imaging support to their omics projects (i.e. OMERO functionality). The *Project Wizard* uses the *OMERO client* library to create the appropriate metadata structure in the OMERO server, respecting the correspondences with metadata entities in the openBIS server. In technical terms, entity correspondence is achieved by providing the OMERO entities with the unique identifiers of the corresponding entities in openBIS.

An additional portlet application was developed to allow easy web-based access to imaging data from qPortal. The *Image Viewer* application (https://github.com/qbicsoftware/omero-client-portlet) uses the *OMERO client* component to query metadata information (e.g. image identifiers, spatial size, image time points and channels) and image thumbnails from the OMERO server, and display this information in a structured manner that follows the proposed metadata model coupling. I.e. a user must select a previously created project and sample (as created by the *Project Wizard*) to access the associated imaging data (**Fig. 4A**). Through the *Image Viewer* application, users can directly access a full, 5D view of any image, which is provided by the OMERO.web server (**Fig. 4B**). To allow direct access to the 5D image viewer, the *Image Viewer* application creates an HTTP session with the OMERO.web server via the OMERO json API, providing the user with a consistent single-sign-on (SSO) experience. Similarly, the OMERO.iviewer application (an OMERO.web plugin) can be employed to facilitate web-based annotation of ROIs (**Fig. 4C**).

**Fig. 3.**
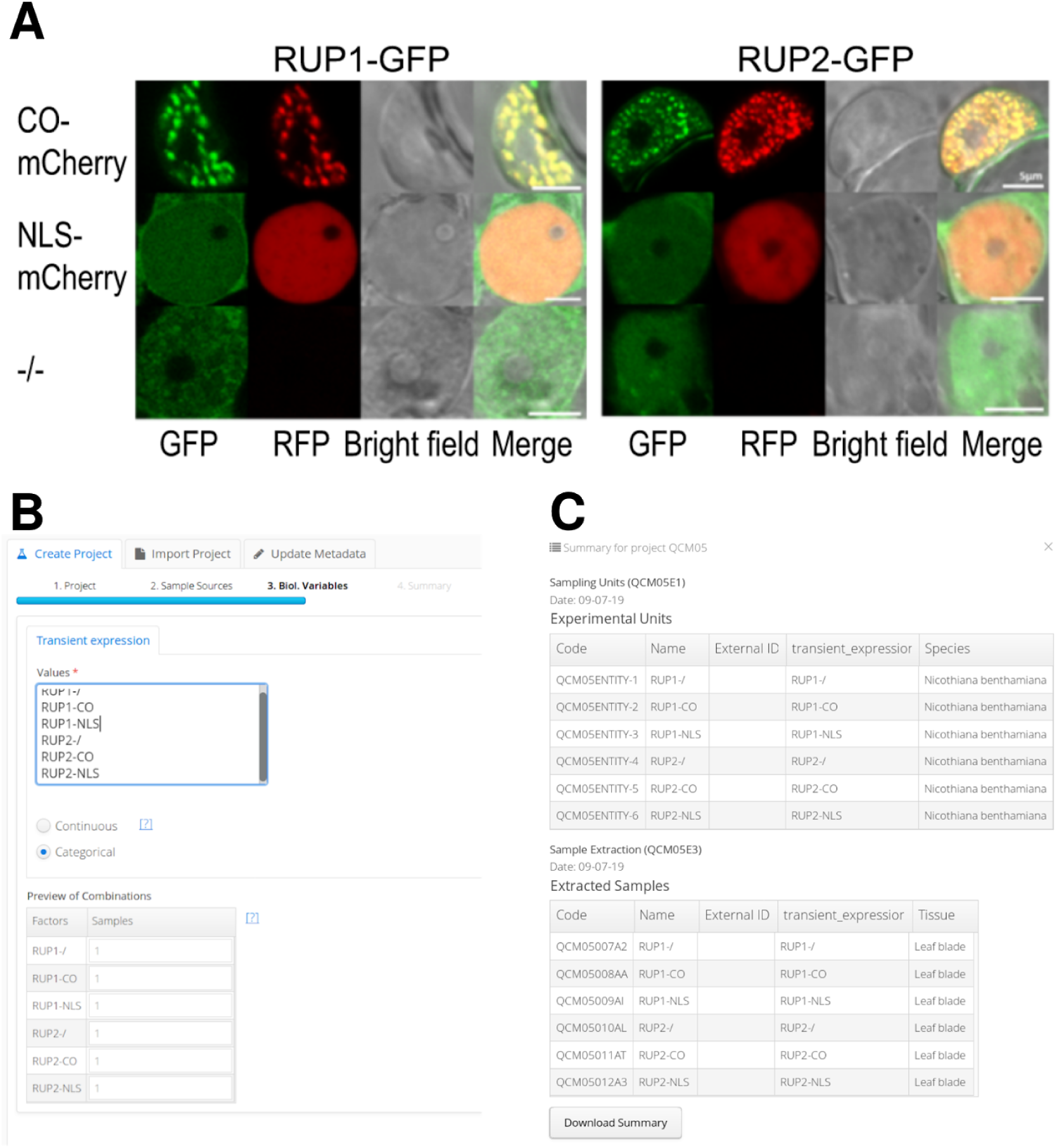
Registration of the Arabidopsis RUP experiment. **(A)** Colocalization experiment of RUP1-GFP and RUP2-GFP and CO-RFP using confocal microscopy data, adapted from (Arongaus et al., 2018). **(B)** The *Project Wizard* application during project creation. **(C)** Summary of the project metadata in the *Project Browser*.

**Fig. 4.**
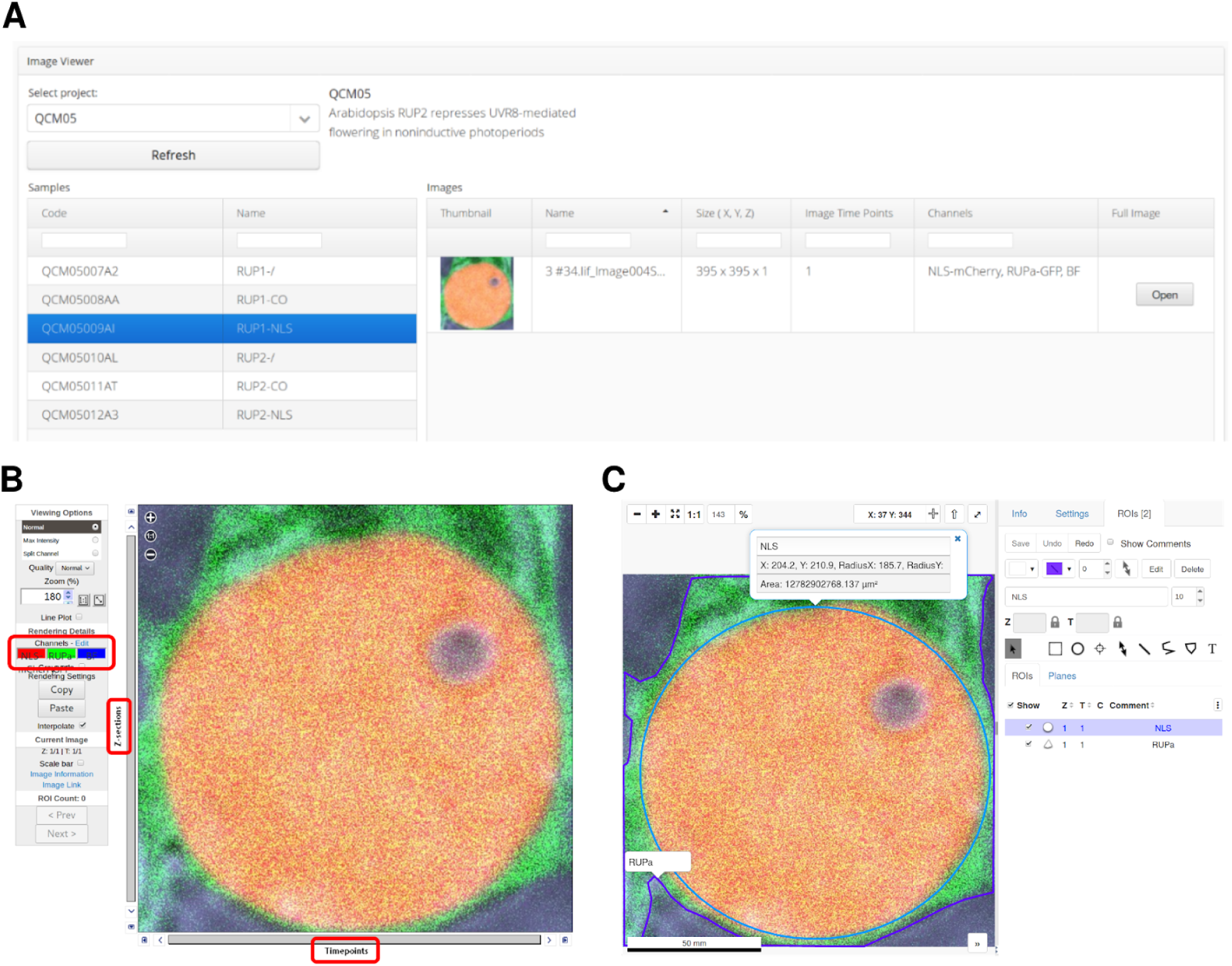
The *Image Viewer* application while accessing the previously uploaded confocal images. **(A)** View of the *Image Viewer* application. **(B)** The 5D image viewer of the OMERO.web server provides an efficient, web-based imaging interface with functionality to visualize 5D images and allows easy navigation in the spatial, temporal and channel dimensions (red boxes). **(C)** The OMERO.iviewer application allows web-based annotation of ROIs, which can be easily stored in the OMERO database.

Imaging data registration is achieved by creating a specialized openBIS *dropbox* and respective ETL procedure, which can connect to the OMERO server through the Python API, and uses the OMERO.importer application as an external tool. In short, the OMERO.importer uploads raw imaging data and uses Bio-formats to extract all metadata from open and proprietary file formats before mapping it to the OMERO metadata model and registering it accordingly. Subsequently, registration of the uploaded images in the openBIS server is achieved by creating specialized metadata entities that contain the unique OMERO identifiers of each uploaded image. Once an image with a correct sample identifier has been uploaded using the ETL routine, the OMERO server will generate a thumbnail of the image and allow access to all associated metadata and raw data via the *Image Viewer* application. **Fig. 2** depicts the activity diagram of the aforementioned ETL routine.

### Use cases

We deployed our newly developed infrastructure for testing with several use cases, ranging from basic single layer experiments to large multi-scale clinical studies. Here we illustrate the broad applicability of an architecture for integrated management of omics and imaging data using two prototypical use cases. The first use case focuses on fluorescence microscopy data from plant biology research, while the second use case deals with a 3-layer omics dataset and multi-modal imaging of liver cancer.

#### Use Case 1: Arabidopsis RUP2

This study aimed to improve our molecular understanding of photoperiodic flowering, a light-controlled mechanism in plants, which allows them to optimize the timing of the transition from vegetative growth to flowering, by responding to seasonal changes in day length (Arongaus et al., 2018). An experiment was designed to test the interaction between RUP1, RUP2 (repression of UV-B photomorphogenesis 1 and 2, respectively) and CO (constans transcription factor), two proteins involved in photoperiodic flowering control, using epidermal leaf cells of *Nicotiana benthamiana* that were transiently transformed. The localization of GFP-tagged RUP1 or RUP2 when coexpressed with either CO-mCherry or NLS-mCherry (nuclear localization signal) was investigated with confocal laser scanning microscopy (**Fig. 3A**).

The fluorescence microscopy data acquired in the abovementioned experiment was registered into our prototype version of qPortal with OMERO support. The initial step in registering this experiment was to create a project using the *Project Wizard* application, and input the experimental design alongside sample information (e.g. species, tissue, experimental condition). The *Project Wizard* will then create the metadata structures in the backend and generate unique identifiers (codes) of each of the biological samples (**Fig. 3B**). Once a project containing the colocalization experiment was created, it could be accessed using the *Project Browser*, an application that can provide a summarized view of the project or display an in-depth diagram representing the metadata structure and entities (**Fig. 3C**). Using the sample codes generated by the *Project Wizard*, confocal images of each of the experimental samples in the colocalization analysis were uploaded using the dropbox mechanism (see implementation). Once all data was uploaded, images could be accessed through the *Image Viewer* application, they were easily findable using sample codes and metadata filters (**Fig. 4A**), and could be visualized using the 5D image viewer of OMERO.web server (**Fig. 4B**).

#### Use Case 2: Hepatocellular Carcinoma Clinical Study

We recently applied our infrastructure to a clinical study involving patients with Hepatocellular Carcinoma (HCC, NCT02372162, “Fingerprint characterization of advanced HCC”). The study aimed at predicting the patient’s response to the drug sorafenib and identifying the molecular or image determinants for therapy response. The disease progression was followed by medical imaging analysis before therapy, after 4 weeks of therapy and 4-6 months after therapy in case of progression. The imaging data was acquired at the Tübingen University Hospital and included magnetic resonance imaging (MRI) combined with Positron-Emission Tomography with two different tracers (PET), and Computerized Tomography (CT) scans for tumor and liver perfusion analysis. CT-guided liver biopsies were additionally acquired. Molecular genetics and gene expression data profiles were obtained from the tumor biopsies and from the healthy liver tissue at the same time points. Next-generation sequencing (NGS) was performed at the medical genetics facility in Tübingen. Metabolite markers in blood and urine of the patients were measured using Nuclear Magnetic Resonance (NMR) at the Werner Siemens Imaging Center, to follow progression markers of the disease and the drug’s pharmacokinetics. The data generated at the three facilities was integrated into our infrastructure (**Fig. 5**).

**Fig. 5.**
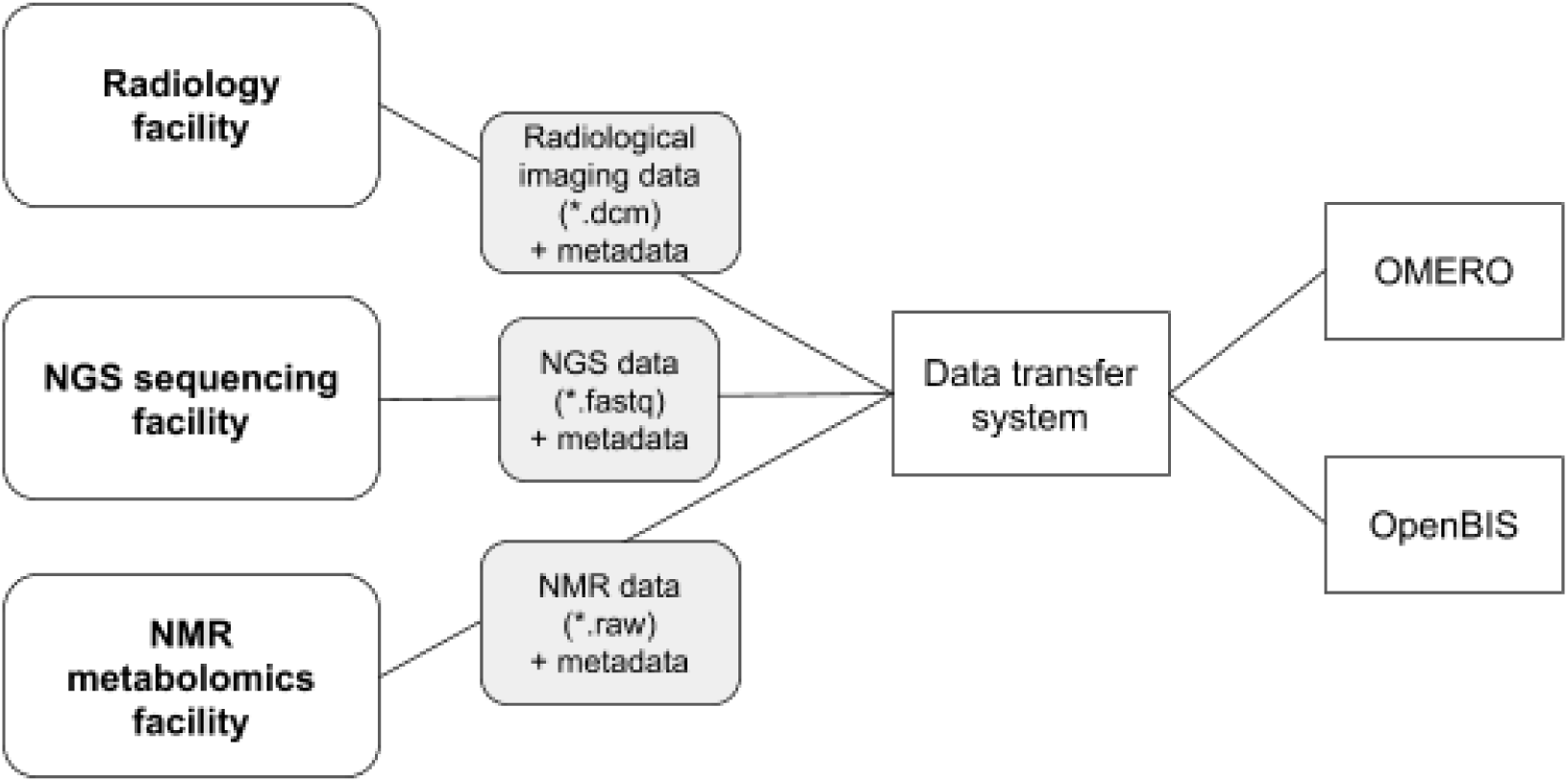
Schematics of the data transfer procedure from the data acquisition facilities. Data and metadata import was achieved via data transfer systems: a Datamover instance in the imaging and medical genetics facilities (openBIS, https://wiki-bsse.ethz.ch/display/DMV/Home), or via the in-house developed dync command line tool (https://github.com/qbicsoftware/dync). The data was then handled on an incoming server in an ETL process and registered in the openBIS and OMERO platforms.

The study generated a total of 3.1 Terabytes of data, comprising NGS (transcriptomics and genomics), radiological imaging, including radiomics analysis, and NMR metabolomics datasets (**Fig. 6A**). The different project partners were given access to the complete study data through the qPortal. The Image Viewer portlet provides an overview of the acquired radiomics data, and enables the display of the radiologist annotations corresponding to the tumor regions (**Fig. 6B and C**). By following the OMERO link, the reconstructed 3D images can be inspected (**Fig. 6C**). Finally, the radiomics findings were contrasted with molecular information about the tumor, provided by the other omics modalities stored in our portal. The genetic variations of the patient’s tumors were analyzed, in order to identify possible driver genes responsible for tumor development and regression (**Fig. 6D**).

**Fig. 6.**
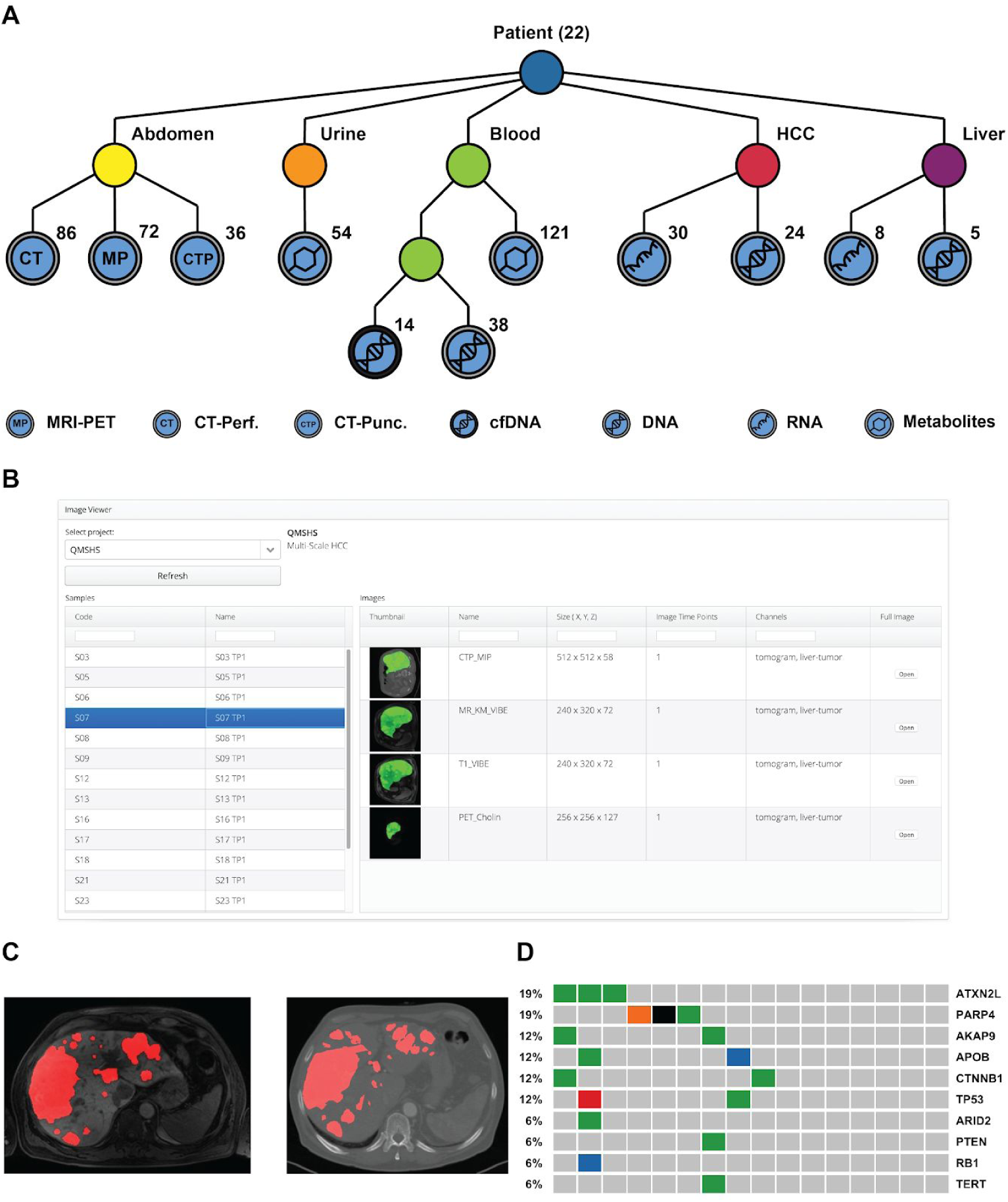
Data availability for the HCC clinical study use case. **(A)** Schematics of the datasets collected for the clinical trial. Radiological imaging data (CT-perfusion (CT), CT-guided biopsy (CTP), MRI-PET (MP) with FDG and Choline tracers) and multi-omics data (metabolomics, genomics and transcriptomics) are integrated into the openBIS metadata model. All data is available to the project partners through our portal. The graph representation allows a quick visualization of the experimental design (Friedrich et al. 2015; Friedrich et al. 2018). **(B)** The image viewer portlet offers an overview of the available imaging data for the project, and provides access to selected metadata. The user is also provided with a link to a full image viewer for the reconstructed tomograms. **(C)** Sample images of MRI (left) and CT (right) of the abdomen for one of the patients. If tumor annotations by the radiologists are available, they can be displayed as an extra channel (here in red). **(D)** Summary of the tumor genomics findings for all patients with available genomics data (N = 16), selected mutated genes are displayed.

## Discussion

We presented a data management architecture for life science research capable of handling projects with complex experimental designs, and managing multi-omics data in conjunction with imaging data from the majority of microscopy and medical imaging disciplines (e.g. PET, electron and fluorescence microscopy). The integration of these datasets was achieved by integrating an OMERO server into qPortal, a web-based multi-omics platform built on openBIS. An important feature of this approach is the integrative metadata model, which results from the coupling of the OMERO and openBIS models. This model defines clear metadata boundaries, assigning project, experimental design and omics metadata to the openBIS server, while using OMERO to store metadata that describes available imaging data and its acquisition process. The objective of these metadata boundaries is to take advantage of the strengths of each platform. However, it is important to carefully assign metadata elements to at least one platform, and avoid replicating metadata on both servers, since this may lead to significant complexity in future metadata curation.

A salient feature of the proposed architecture is the SOA-based approach used to implement a loose-coupling between openBIS and OMERO, as modular software components of the qPortal backend. In particular, for defining the metadata connections between the qPortal and OMERO models, which benefited from similarities in the main structure of the models (see **Fig. S1**), and allowed their complementary characteristics to be leveraged with few restrictions. Specifically, the OMERO server was treated as a modular component, responsible for storing raw imaging data and metadata related only to imaging datasets and their acquisition process, thereby leveraging OMERO’s core functionality while allowing qPortal and openBIS to continue to enforce FAIR data management, and to store omics data and metadata related to the experimental design of projects.

In SOA, a variety of specialized and modular components provide services via APIs, allowing them to operate in concert within a distributed system. In this context, qPortal can serve as a suitable distributed framework for data management of multi-omics and imaging data, since it provides additional abstraction layers where the middleware needed to implement the communication protocol between the OMERO and openBIS components can be developed. Moreover, qPortal operates as a flexible, web-based platform with a set of applications for scientific data management, and capable of deploying custom Java applications for a particular omics or imaging modality (e.g the *Image Viewer* application).

An alternative approach to the proposed architecture would be to extend a single platform (e.g. openBIS or OMERO), leading to a monolithic backend system with aggregated functionality. This type of information systems tend to be highly complex, lacking flexibility and scalability. On the one hand, when extending qPortal or openBIS to accommodate biomedical imaging data, it is relevant to notice that while qPortal is capable of FAIR management of complex multi-omics data, it was not designed to support raw imaging data and metadata from a large variety of microscopy and medical imaging disciplines without a significant extension to its core functionality, that would likely lead to replication of most of the functionality and the metadata model offered by OMERO, e.g. an image rendering engine, proprietary file format parsers, and database support for ROIs (see the OMERO platform section).

On the other hand, while OMERO offers an extensible metadata model and allows customization to support omics data, as demonstrated by an extension to accommodate data from a Genome Wide Association Study (GWAS) of human autoimmune diseases (Sanna et al. 2010; Allan et al. 2012), it is still unclear if complex and highly heterogeneous multi-omics experiments can be efficiently stored and queried using the OMERO server alone. Extending the OMERO server to efficiently enforce FAIR-compliant, multi-factorial experimental designs, storage and annotation of multiple, high-throughput omics disciplines, would likewise require significant software development and long-term support, that would also replicate functionality already offered by openBIS and qPortal.

Therefore we opted for a SOA approach to efficiently integrate management of omics and biomedical imaging data, allowing us to leverage several technologies in a modular way. By abstracting the openBIS and OMERO servers as backend service components within a distributed system (qPortal), it was possible to selectively use the metadata and raw data storage facilities of each component during the execution of the required data management operations, while maintaining the integrity of metadata structures across all backend components.

We have also presented two use cases where qPortal with an integrated OMERO backend was used to efficiently store imaging data in a multi-omics environment. The Arabidopsis RUP use case described the registration process of a plant biology project containing data from a confocal scanning microscope, and showed how such imaging data could be easily accessed via the web interface of qPortal. The second use case presented a hepatocellular carcinoma study with highly heterogeneous medical imaging and multi-omics data, allowing physicians to relate several disciplines (e.g. NGS, gene expression data, confocal, PET and MRI imaging data) via the metadata structure of the project, which relates biological samples entities (and all their associated datasets) with the corresponding patient and time point of the clinical study.

Our data management architecture allows the deployment of highly scalable data repositories capable of linking omics and biomedical imaging modalities. Leveraging the interoperability of backend components and storing metadata in an integrative and standardized manner, allows searching across studies to derive datasets of integrated omics and biomedical imaging data, thus facilitating multi-modal data reusability. Such datasets are particularly attractive given the complementary nature of omics and biomedical imaging modalities, since omics disciplines enable the simultaneous measurement of a large variety of molecular components and species *in vitro*, but lack the spatio-temporal information provided by imaging modalities. Moreover, since significant breakthrough in microscopy techniques permit the visualization of cellular structures *in situ*, at ever increasing resolutions (e.g. super-resolution light microscopy (Sigal et al. 2018), direct electron detectors (Cheng 2015)), and both omics and microscopy modalities have reached single-cell sensitivity, the need to correlate such datasets has significantly increased (Hériché et al. 2019).

Joint analysis of imaging and omics data can provide new insights into biological processes. Multi-modal data analysis can be accomplished by independently analyzing data from different modalities, that has been extracted from the same biological sample, ideally using reproducible bioinformatics workflows (Ewels et al. 2020). Integrated multi-modal analysis is also possible by using higher-order data representations or by employing deep learning methods (Hériché et al. 2019). Importantly, supervised machine learning methods, such as deep neural networks, rely on the availability of large training datasets annotated with rich and high quality phenotypic metadata, underscoring the importance of multi-modal and FAIR compliant data repositories.

## Conclusion

Here we present a SOA-based method to integrate an image management system, the OMERO server, as a modular, backend component of qPortal, to allow the integrated management and analysis of multi-omics and biomedical imaging data. The implementation of a structural coupling between the openBIS and OMERO metadata models, the development of software components to facilitate the communication with the OMERO server, and an extension to the data management operations of qPortal, facilitated storage and analysis of raw data and metadata from various omics, microscopy and biomedical imaging modalities in an integrative manner, with the ability of accepting metadata queries from web-based, scientific applications. The applicability of the proposed architecture was demonstrated in two use cases, a plant biology study using confocal scanning microscopy, and a clinical study on hepatocellular carcinoma, with data from heterogeneous medical imaging and omics modalities.

As emerging developments in omics and biomedical imaging drive the increase in resolution, modality, and throughput of data generation in life science studies, following SOA principles in extending, integrating, or building the required information systems for long-term storage and FAIR management of these valuable digital assets, offers a flexible and scalable approach to manage a variety of heterogeneous, multi-omics and imaging data repositories.

Such FAIR and scalable data management infrastructures for data repositories, capable of tackling the ever increasing volume and complexity of omics and biomedical will not only enable high-throughput, multi-modal data management in life science, but also allow the generation of large and highly multi-dimensional datasets, via metadata queries across stored scientific projects, enabling data-driven analysis and training of machine learning models in future predictive applications.

## Acknowledgements

We acknowledge funding from BMBF MultiscaleHCC, DFG SFB/TR 209, DFG SFB 1101, DFG SFB/TR 261, Excellence cluster microbiology. SN and LKC acknowledge funding from SFB 261 and FW, SzO-K. LKC and KH acknowledge funding from SFB 1101 (projects D02 and Z02). GG acknowledges funding from the German Ministry of Research and Education (BMBF, grant no. 01ZX1301F).

SN acknowledges funding from Deutsche Forschungsgemeinschaft (core facilities initiative, KO-2313/6-1 and KO-2313-2, Institutional Strategy of the University of Tübingen, ZUK 63). Furthermore, SN acknowledges funding by the Sonderforschungsbereich SFB/TR 209 “Liver cancer” of the Deutsche Forschungsgemeinschaft (DFG), as well as from the Deutsche Forschungsgemeinschaft (DFG, German Research Foundation)-Project-ID 398967434 - TRR 261 and the Exzellenzclusters “Controlling microbes to fight infection” (CMFI), EXC-2124.

The clinical study (NCT02372162) was funded by the German Ministry for Education and Research (BMBF, eMed/Multiscale HCC, FKZ 01ZX1301A und 01ZX1601G, N.P.M., M.B.).

## Supplementary Material

**Fig. S1.**
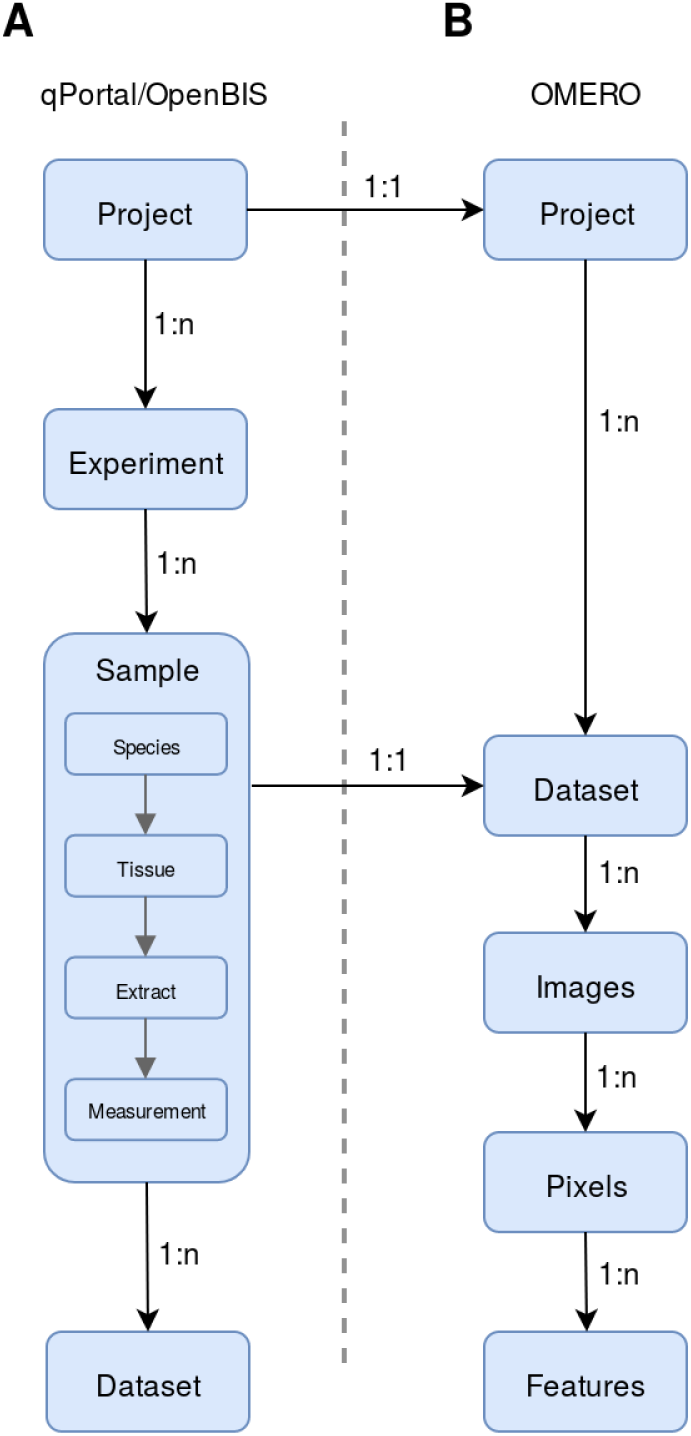
Diagram of the unified metadata model, depicting the coupling between the underlying models and the cardinality relationship between metadata entities. **(A)** The hierarchical metadata model used by qPortal to describe the experimental design of a research project, containing general information of sample biology. **(B)** The hierarchical metadata model used by OMERO, focused on describing imaging data.

## Notes

### Competing Interest Statement

The authors have declared no competing interest.

